# The major pathogen *Haemophilus influenzae* experiences pervasive recombination and purifying selection at local and global scales

**DOI:** 10.1101/2024.10.16.618562

**Authors:** Neil MacAlasdair, Anna K. Pöntinen, Clare Ling, Sudaraka Mallawaarachchi, Janjira Thaipadungpanit, Francois H. Nosten, Claudia Turner, Stephen D. Bentley, Nicholas J. Croucher, Paul Turner, Jukka Corander

## Abstract

*Haemophilus influenzae* is a major opportunistic human pathogen which causes both non-invasive and invasive disease. The *H. influenzae* type b (Hib) vaccine has led to a significant reduction of invasive Hib disease, but offers no protection against colonisation or disease by unencapsulated non-typeables (NT) or non-b serotypes, and *H. influenzae* remains a public health burden worldwide, with increasing reports of multi-drug resistance (MDR). Despite this, there is no comprehensive understanding of the species’ global population structure. To advance understanding about the evolution and epidemiology of the species, we whole-genome sequenced 4,475 isolates of *H. influenzae* from an unvaccinated paediatric carriage and pneumonia cohort from northwestern Thailand. Despite no Hib immunisation, serotype b was uncommonly found (5.7%), while 91.7% of isolates were NT. We identified a large number of nearly pan-resistant lineages that were mostly NT, and discovered that no lineages were enriched among disease samples, suggesting the ability to cause invasive disease is not restricted to any subpopulation of the species. Extensive population genetic analyses of our data combined with a worldwide collection of 5,976 published genomes revealed a highly admixed population structure, low core genome nucleotide diversity, and evidence of pervasive negative selection. The combined data confirm that MDR lineages are not confined to our cohort, and their establishment globally is an urgent concern.

## Introduction

The nasopharynx is the natural habitat of the bacterium *Haemophilus influenzae*, where it exists in asymptomatic carriage, while frequently translocating to other body sites such as inner ears, lungs, and sinuses, causing a range of disease manifestations ^1^. Most common of these is acute otitis media (AOM), which is one of the leading causes of antibiotic prescriptions in children. Global estimates suggest over 700 million AOM cases per annum caused in total by any bacterial pathogen and a significant fraction of these lead to further complications and sequelae, particularly in low- and middle-income countries (LMICs) ^2^. After licensing of the polysaccharide-protein conjugate *H. influenzae* type b (Hib) vaccine in the late 1980s, its adoption into national vaccination programs worldwide has led to a significant reduction of Hib colonisation and its associated invasive disease manifestations, such as meningitis and pneumonia. However, the vaccine does not protect against colonisation by other serotypes or unencapsulated non-typeable *H. influenzae* (NTHi). Therefore, *H. influenzae* remains a significant cause of AOM, sinusitis, conjunctivitis and pneumonia, and consequently is an important public health burden globally. A particular concern has arisen from the widespread antibiotic resistance observed in some strains of NTHi ^3^.

The evidence for NTHi as an important cause of paediatric community-acquired pneumonia (CAP) has been summarised in comprehensive reviews ^4,5^. Determination of aetiology in paediatric CAP remains a challenge, with a minority of cases being bacteraemic. However, specimens obtained via bronchoscopy revealed NTHi to be the dominant bacterial pathogen in 250 Belgian children with recurrent or non-resolving CAP ^6^. Nasopharyngeal colonisation by non-Hib / NTHi, especially at higher densities, has also been shown to be associated with paediatric CAP in LMICs ^7,8^.

Whole-genome sequencing studies of *H. influenzae* have been mainly conducted from smaller-scale collections of disease cases ^9,10^, but rarely from large-scale collections of both carriage and disease isolates of the same population. Furthermore, few studies have been conducted in LMICs, where nasopharyngeal pathogen colonisation rates and the burden of CAP are generally much higher than in high income countries. As a consequence, the genetic population structure and evolutionary dynamics of the species remain poorly understood in LMIC settings and at a global scale ^11^.

This motivated us to conduct a longitudinal paediatric cohort study of both healthy colonisation and pneumonia among a large birth cohort in a population located in Northwestern Thailand, the Maela camp for displaced persons. The densely populated camp is located on the Thailand-Myanmar border and provided a unique opportunity to systematically sample both carriage and disease cases in a pre-Hib vaccine population. Here we detail results from the whole-genome sequencing (WGS) of isolates from our longitudinal cohort study, and also of further analyses performed on the Maela data combined with all publicly available high-quality *H. influenzae* WGS data with known year and geographical location of isolation. This combined collection of 9,849 genomes allowed us to conduct genomic analyses of the species at an unprecedented global scale and sampling intensity, and provided novel insight into how its high levels of recombination shape its global population structure.

## Results

### Serotype distribution across the Maela paediatric population

Infants in the Maela cohort carrying *H. influenzae* (Fig. 1A, B) were predominantly colonised by NT *H. influenzae*, despite lacking immunisation against Hib (Table 1). Out of 3,970 isolates that passed final QC filters, the counts and estimated frequencies of the six different serotypes (Watts and Holt, 2019) and non-typeable (unencapsulated) isolates are listed both with and without host deduplication in Table 1. Notably, non-typeable isolates made up 91.7% of all isolates, and serotype b isolates are the second most prevalent, making up 5.7% of the population. The remaining 5 serotypes account for less than 1% of the population each. Serotypable isolates generally form monophyletic lineages on the tree (Fig. 2). Two isolates, one NT and one serogroup b by agglutination, gave only partial *in silico* capsule typing results due being unable to identify the entire capsule locus in the data. *In silico* capsule typing was generally congruent (overall congruence 95.3%) with the agglutination-based phenotypic serotyping (except for serotypes d and e), and was corrected by the latter in those 28 cases where the serological typing indicated a serotype for a non-typeable *in silico* type. These included serotype a (n = 1), b (n =19), d (n =3), and e (n = 5). Our results are reasonably well in line with earlier comparisons between agglutination-based and *in silico* typing (98-100% congruence)^12,13^, however, the higher level of discrepancy observed here could be due to the much larger and more diverse set of genomes considered.

**Figure 1.**
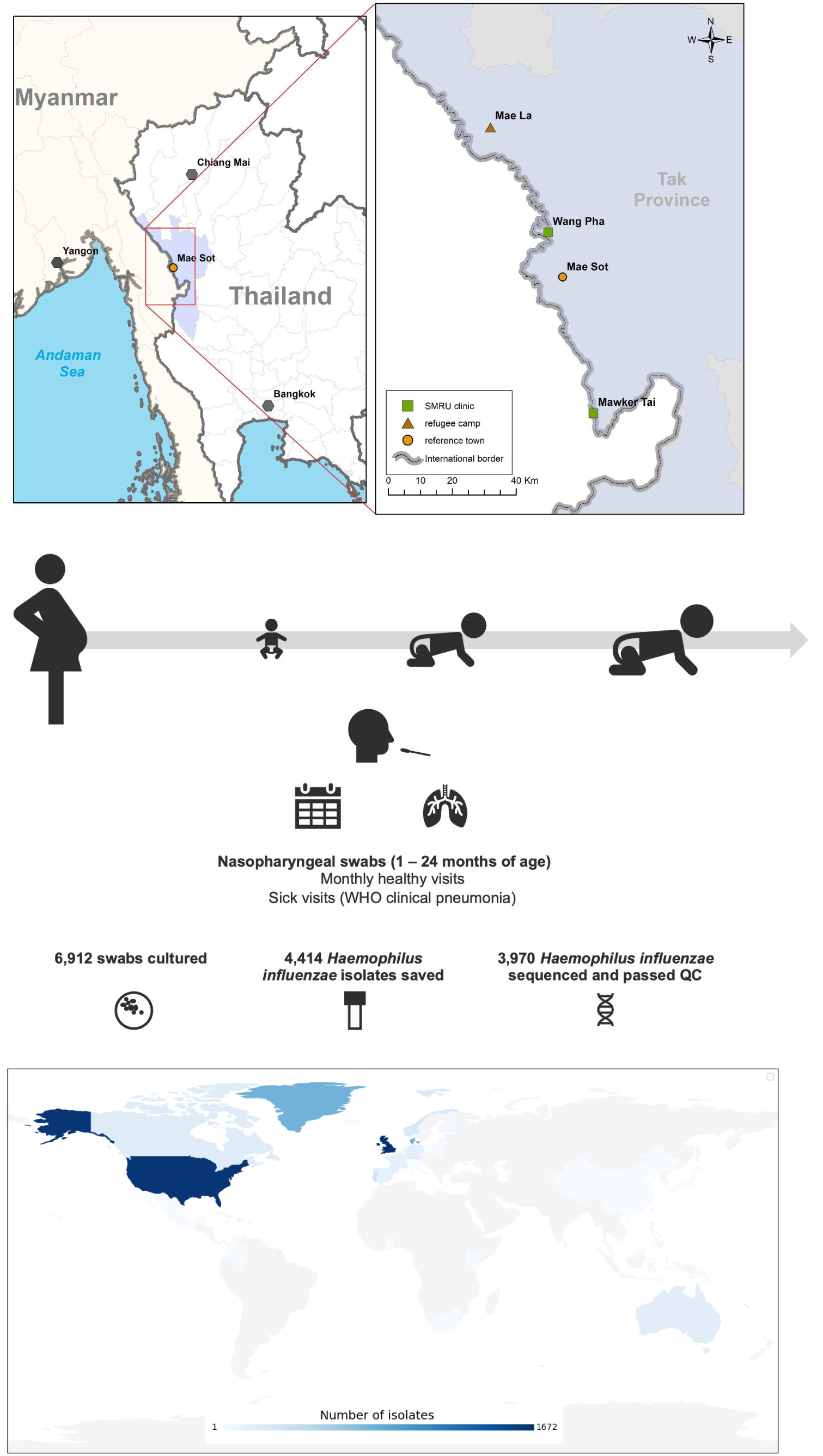
A: Geographical location of the study site. B: Cohort design and sample processing. C: Global map coloured by number of isolates per country of origin in the systematic public collection of global *H. influenzae* isolates.

**Figure 2.**
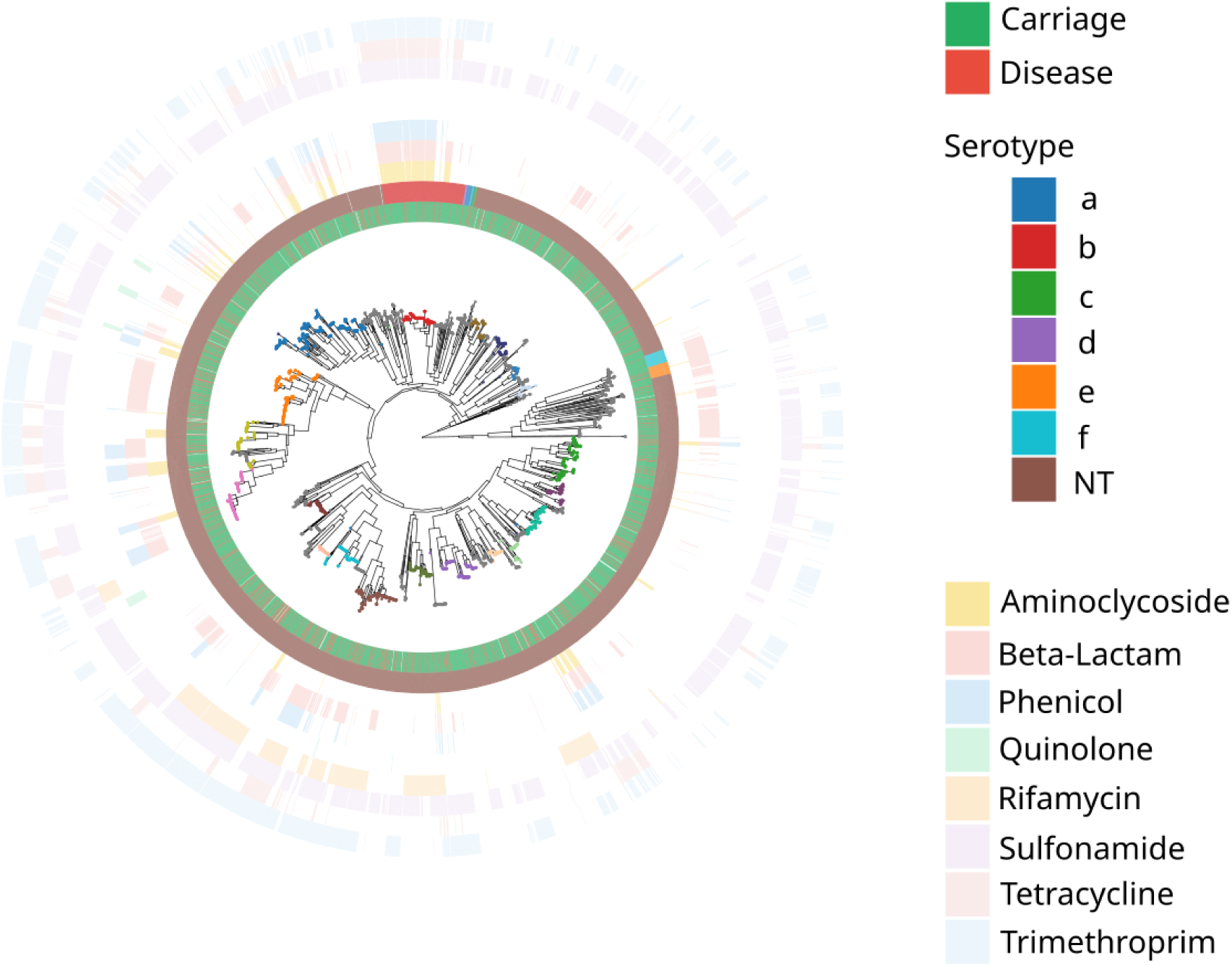
Phylogeny of Maela *H. influenzae* genomes for 3,970 isolates from carriage and pneumonia samples (inner ring), estimated using FastTree v.2.1.10 on the core-genome alignment mapped against the *H. influenzae* reference 86-028NP (NC_007146.2). The 20 largest PopPUNK clusters (>50 isolates) are indicated by coloured dots at the tips of the phylogeny, while smaller clusters are grey. *In silico* serotypes (second ring), inferred by using Hicap v.1.0.3, and AMR profiles (eight outer rings), screened with AMRFinderPlus v.4.0.3, are shown by colour as indicated in the legend. AMR = antimicrobial resistance. An interactive online phylogeny, with additional metadata including cgMLST and cgMLST Cluster data is available at the following link: https://microreact.org/project/oMm8PFCoG2429JwiDBpdru-maela-h-influenzae

**Table 1.**
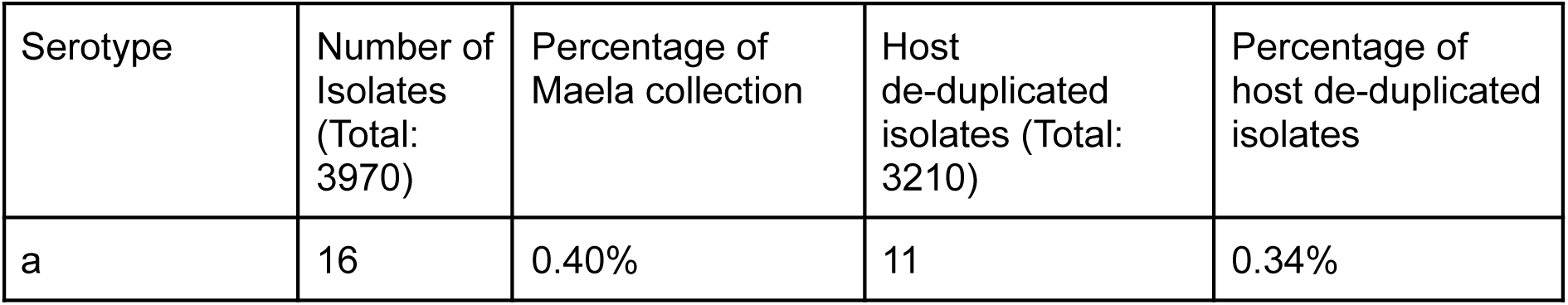
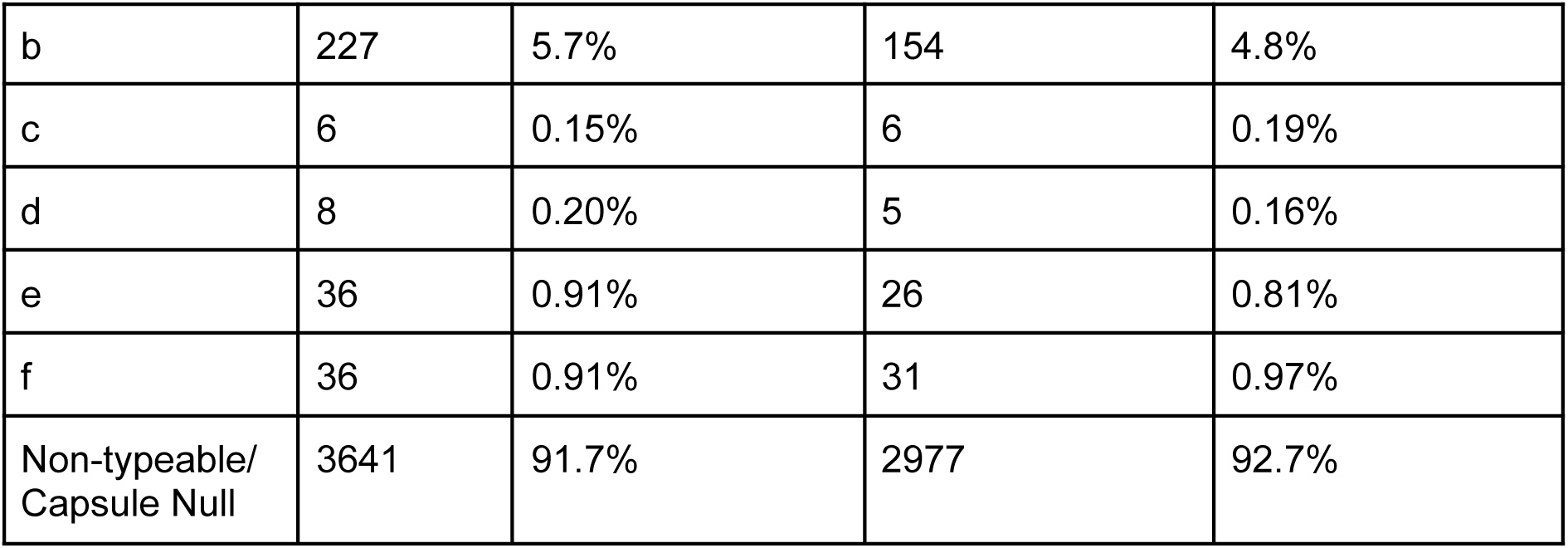
Proportion and counts of the number of isolates of each serotype and non-typable/capsule null isolates. Serotypes were determined with both agglutination and *in silico* using sequence data (Methods). Due to the longitudinal nature of the sampling, we also removed isolates which were collected from the same host on consecutive sampling times and were likely to be clonally related (Methods).

### Genetic population structure of *H. influenzae* in Maela

Core genome MLST (cgMLST), has previously been used to study *H. influenzae^11^*, but when we applied it to the entire global dataset (see **Population genetic analyses of the global dataset**), neither the typing nor clustering by allelic profiles was able to provide meaningful insight into the population structure of *H. influenzae* in the Maela cohort, as the diversity in the allelic profiles was too high. Of note, the number of cgMLST allelic profiles and overall nucleotide diversity are not always strongly correlated, since it is possible for a large number of very low-frequency mutations to generate a large number of allelic profiles (and allelic mismatches), even though nucleotide diversity across the pangenome remains low. Using PopPUNK to cluster the Maela isolates was largely able to identify monophyletic clusters (Fig. 2), however, PopPUNK’s final, optimal clustering divided the population into many small clusters. The largest PopPUNK cluster contained 349 isolates, with only 13 PopPUNK clusters consisting of at least 100 isolates (out of a total of 122 clusters), and 20 clusters consisting of 50 or more isolates. A large proportion of clusters (50%) contained 10 or fewer isolates. NT *H. influenzae* were observed as the dominating type in both healthy carriage and pneumonia time point samples (Fig. 2), and no particular genetic lineage was overrepresented in pneumonia samples (Fisher’s exact test p-value 0.091, Methods).

### Distribution of AMR determinants

AMR determinants were frequently identified across the phylogeny and strongly associated with the MDR lineages (Methods, Fig. 2). Only one of these MDR lineages was clearly associated with serotype b (Fig. 2), while the remainder consisted of non-typeable isolates. A total of 41 PopPUNK clusters contained at least one MDR isolate, indicating repeated acquisition of AMR determinants across the population.

In the Maela host-deduplicated dataset, most of the MDR isolates (resistance against at least four out of nine antibiotic classes, Methods) (507/3210), were NT (77.3%, 392/507), followed by serotype b (22.3%, 113/507), and two serotype e isolates. Hence, serotype b was clearly overrepresented among the more resistant isolates (overall frequency 4.8%, Table 1), while there were less NT (overall frequency 92.7%, Table 1). Within pneumonia cases (523/3210), 17.0% (89/523) of isolates were MDR, of which 76 were NT, 12 of serotype b and one was serotype e. Within non-pneumonia cases (2687/3210), 15.6% (418/2687) were MDR, of which 316 NT, 101 serotype b and one was serotype e, hence the frequency of MDR phenotype was highly similar between the two sample types.

### Quantification of homologous recombination

Since the acquisition of AMR determinants is likely aided by horizontal gene transfer in this naturally transforming species, we quantified the extent of homologous recombination. Mapping Illumina reads from isolates in the same PopPUNK cluster against long-read reference assemblies to produce whole-genome pseudoalignments, to be used as inputs to SNP-density based recombination analysis (Gubbins) was not a feasible approach to quantify recombination in the entire Maela cohort due to the size of the dataset and the large number of PopPUNK clusters present.

Consequently, we leveraged the aligned pangenome genes for the 3,970 Maela isolates to perform per-gene recombination inference (Methods). Of the pangenome of 7,015 genes, at least one recombination event between PopPUNK lineages was identified in 2,672 genes (38%). On average, 193.36 recombination events were identified per gene (including recombination-free genes) and the frequency of recombination events was significantly correlated with the estimated nucleotide diversity per gene (Spearman’s r = 0.49, *p* < 7·15 × 10^-293^). A substantial proportion of genes with no detected recombination (64.0%) also had zero nucleotide diversity (Fig. 4A). Finally, we also quantified the rate of decay of linkage disequilibrium (LD) in the core genome and compared this with several other common bacterial pathogens analysed in ^14^. This showed that the decoupling of SNPs as a function of base pair distance happens fastest in *H. influenzae* (Fig. 4B), and the rate is considerably elevated compared with other species known to routinely engage in homologous recombination, such as *Campylobacter jejuni* and *Enterococcus faecalis*. Taken together, these results suggest that the *H. influenzae* population within Maela is extremely recombinant, to the extent that it likely reduces the overall level of diversity within the population.

**Figure 3.**
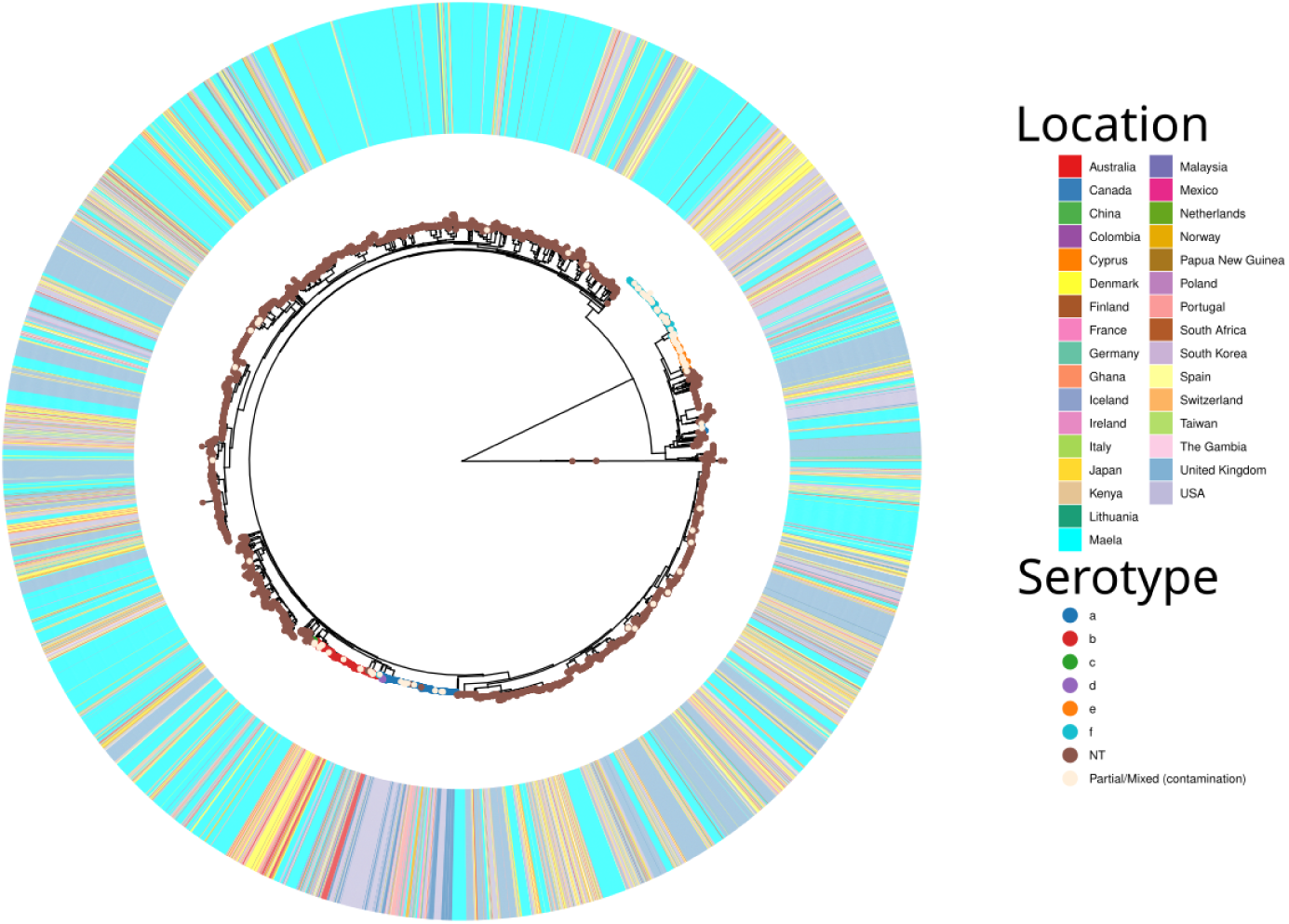
Maximum-likelihood core genome phylogeny of 9,849 *H. influenzae* isolates, combining the Maela cohort and a systematically identified collection of published isolates from around the globe. The phylogeny was estimated using core-genome distances inferred with PopPUNK v.2.4.0. *In silico* serotypes are indicated by the circles on the tips of the phylogeny, isolation location is indicated on the inner ring, as shown by colour as indicated in the legend. An interactive online phylogeny, with additional metadata including cgMLST, cgMLST Clusters, and partial disease state data is available at the following link: https://microreact.org/project/ioyt4oJRSJgeFGK9KmFyVk-global-h-influenzae-core-tree

**Figure 4:**
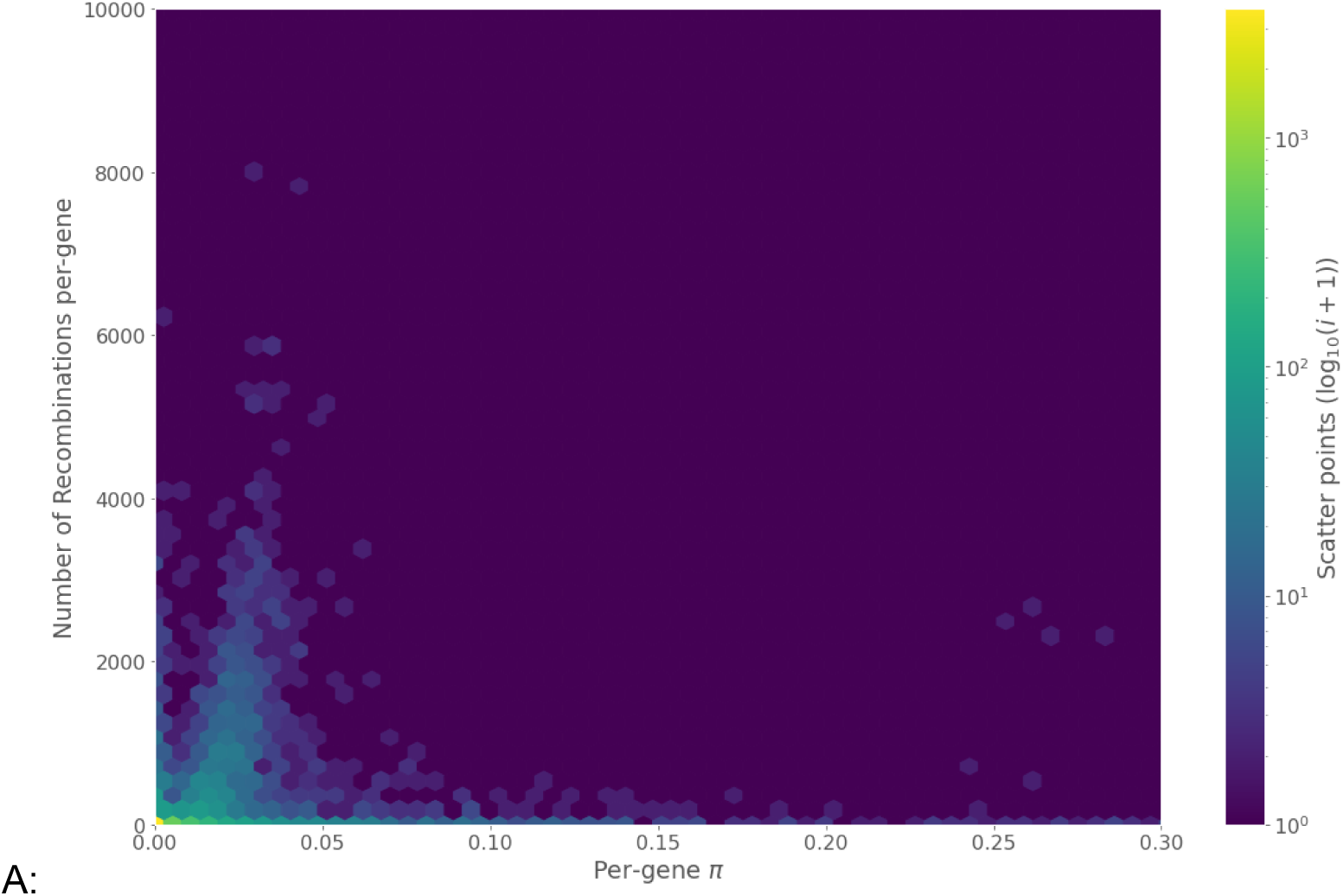
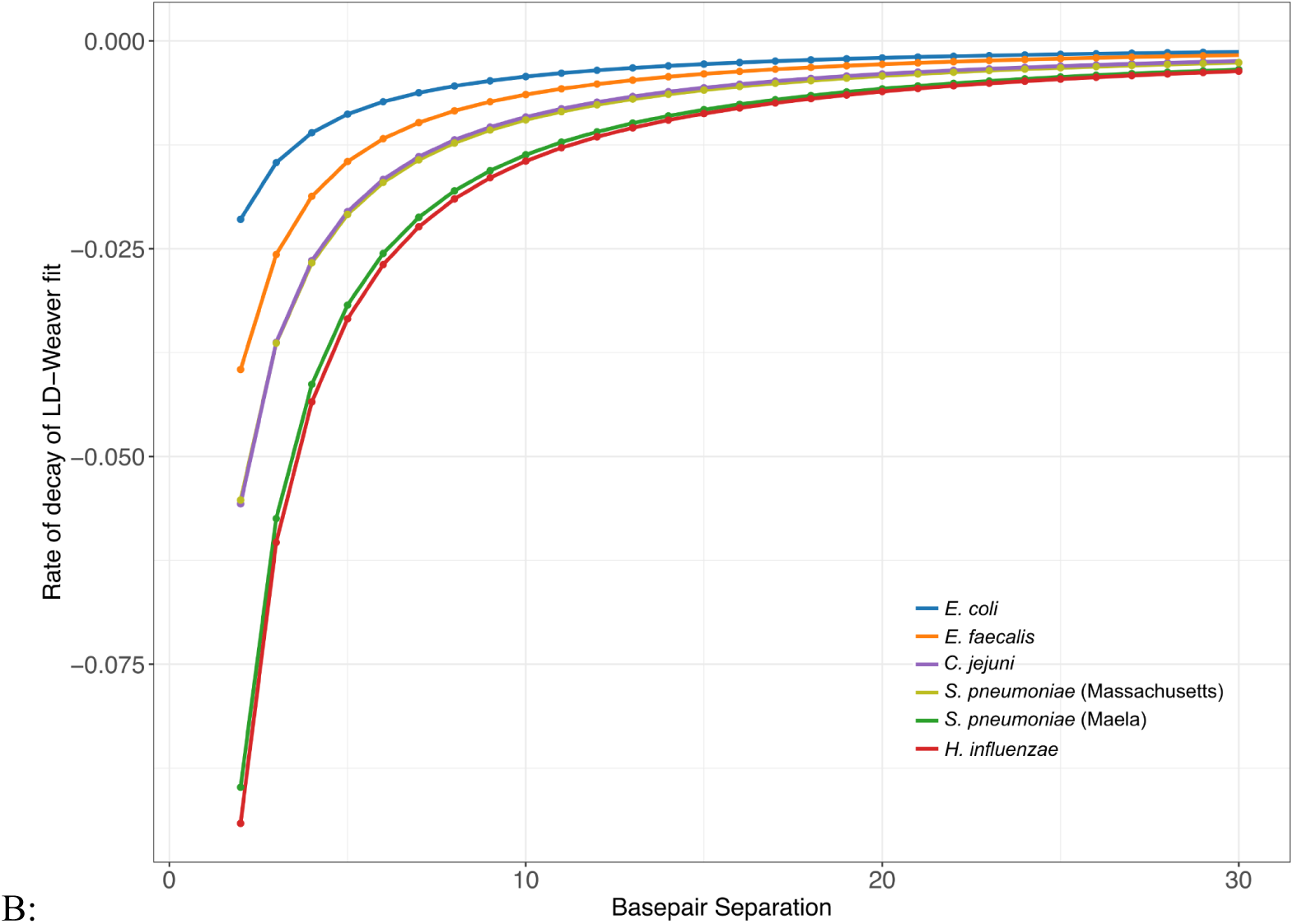
A: Log-scaled hexagon-density scatter plot of the estimated per-gene nucleotide diversity, π, versus the number of recombination events inferred per-gene. Hexagon colours indicate the number of scatter plot points (log-scaled) present within each hexagon. B: The estimated rate of decay in linkage disequilibrium (LD) for the *H. influenzae* Maela cohort, compared with estimated rates for the four species with LD decay functions fitted in ^14^. The shown curves correspond to gradients of the LD decay function, with smaller values indicating faster decoupling of SNPs as a function of distance in base pairs.

### Population genetic analyses of the global dataset

To understand how the genetic variation observed in the Maela cohort samples relates to internationally circulating *H. influenzae*, we combined the study data with a systematic collection of all publicly available *H. influenzae* genome data with basic metadata available (country and year of collection), for a total dataset comprising 9,849 isolates (Fig. 1C, Methods). PopPUNK clustering of the combined dataset successfully identified 752 monophyletic lineages, with the largest cluster composed of a lineage of 483 isolates (>99% serotype a), 20 clusters containing at least 100 isolates (including one cluster each of predominantly serotype a, f, b, and e isolates, remainder NT), and 595 clusters fewer than 10 isolates (81.54% isolates NT). Many larger clusters were paraphyletic according to the core genome tree, while the monophyletic lineages corresponded to small or singleton clusters, likely reflecting a change in the accessory genome of the smaller cluster which had brought the pairwise distances above PopPUNK’s clustering threshold.

The core genome phylogeny of the combined collections (Figure 3, Maela isolates in light blue) clearly demonstrates that the Maela isolates are extensively interspersed within the global population of the species, suggesting rapid cross-border and intercontinental transmission of *H. influenzae*. Furthermore, isolates spanning the sampling window, from 1962-2023 are distributed across the phylogeny and do not form monophyletic lineages made up of temporally restricted isolates. Together, these patterns strongly suggest that the history of migration within the global *H. influenzae* population is sufficiently frequent and extensive to overwhelm any phylogeographical signal of local clonal expansion of lineages.

Due to the extremely low level of nucleotide diversity evident during the recombination analysis of the Maela cohort, we further investigated the overall level of nucleotide diversity across the aligned pangenome of the entire global collection, which consisted of 18,265 genes. Although nucleotide diversity in both core (*n*=1103) and non-singleton accessory (*n*=8843) genes have overlapping ranges, (Figure 5A), core genes are on average significantly less diverse than accessory genes (Two-tailed Mann-Whitney *U* test, *p*=1.205 × 10^-5^, Figure 5A).

**Figure 5:**
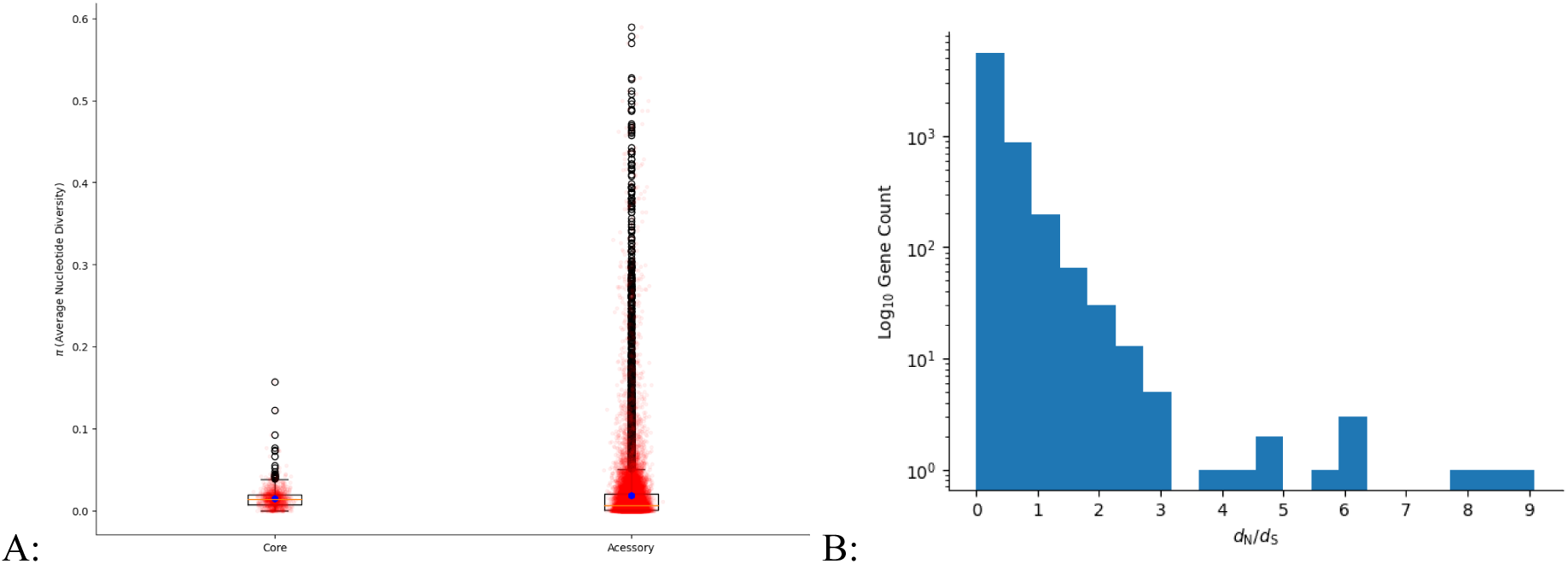
A: Boxplots of the estimated average pairwise nucleotide diversity, π, in each gene of the aligned pan-genome of the combined dataset, split into genes present in 80% or more isolates (core), and genes in less than 80% of isolates (accessory). Blue hexagons indicate gene-frequency weighted average nucleotide diversity across all genes, the yellow line the median, the outer edges the first and third quartiles, and the whiskers 1.5 times the interquartile range beyond those values. Black points indicate outliers, and all data points are plotted in transparent red. B: Log-scaled histogram of the estimated *dN/dS* values, the ratio of nonsynonymous to synonymous mutations, across 6,853 aligned genes from the pan-genome of the combined dataset.

To understand how selective forces may be influencing the diversity observed within the pangenome, we further estimated *dN*/*dS*, the ratio of nonsynonymous to synonymous nucleotide mutations within every gene of the pangenome (Methods). Consistent with the low level of diversity observed, the average *dN*/*dS* value was 0.28, and 96% of the 6,853 genes for which it was successfully estimated (Methods) had *dN*/*dS* < 1 (Figure 5B). This implies that negative selection is widespread across the coding regions of the *H. influenzae* genome. Of the remaining 256 genes (4%) with a *dN*/*dS* estimate, 45 had *dN*/*dS* >2, indicating potential positive directional selection. Further analysis of these genes was undertaken using three statistical tests implemented in the HYPHY v.2.5.60 (Methods), and a few accessory genes possessed extremely strong evidence of selection, where at least two of the three statistical tests rejected the null hypothesis of neutral evolution (Methods). The genes involved included an unnamed gluconate transporter, the BrnT toxin protein, and a third small protein of unknown function. The results of these analyses are illustrated in Extended Data figure 1, 2, and briefly summarised as follows. The unnamed gluconate transporter showed statistically significant results in all three HYPHY tests used, and these tests indicated a branch of the gene phylogeny containing eight isolates, and a specific codon (185) in the protein alignment which have been positively selected for. This branch consists of seven Maela isolates and one isolate from elsewhere, which all possess a structural variant of the unnamed gluconate transporter with a large deletion of a transmembrane domain. The *brnT* toxin gene, the toxin from the BrnT/BrnA type II toxin-antitoxin system ^15^ also showed statistically significant results in all three HYPHY tests, which indicated that a branch of gene phylogeny consisting of two Maela isolates with a large deletion of an alpha helix had recently been subject to positive selection. Finally, the protein of unknown function showed a statistically significant result in two of the three HYPHY tests, identifying a glutamine/valine variable site, with the valine variant primarily associated with Maela isolates, and the glutamine variant primarily associated with isolates from elsewhere. All three of these proteins correspond to low-frequency accessory genes which are globally distributed. Notably, the variants identified as under selection in the *brnT* toxin and the unnamed gluconate transporter are either unique or much more prevalent among Maela isolates, with a similar split association between the two variants of the unnamed protein. This suggests that either the intensive longitudinal sampling frame or the circumstances of the Maela camp may be resulting in elevated statistical power to detect selection, or genuinely stronger positive selection and rapid local adaptation.

Finally, to explore the geographic distribution of MDR lineages in greater detail, we focused on the lineages with at least 50 isolates and 30% resistance prevalence for at least 4 of 9 antibiotic classes. This revealed that all large MDR lineages (n = 3) are widely disseminated internationally, i.e. observed in at least 11 different locations (Fig. 6). Only one of these lineages was dominated by Hib strains and showed evidence of independent capsule switches to serotype a (Fig. 6). The others were composed of non-typeable isolates.

**Figure 6.**
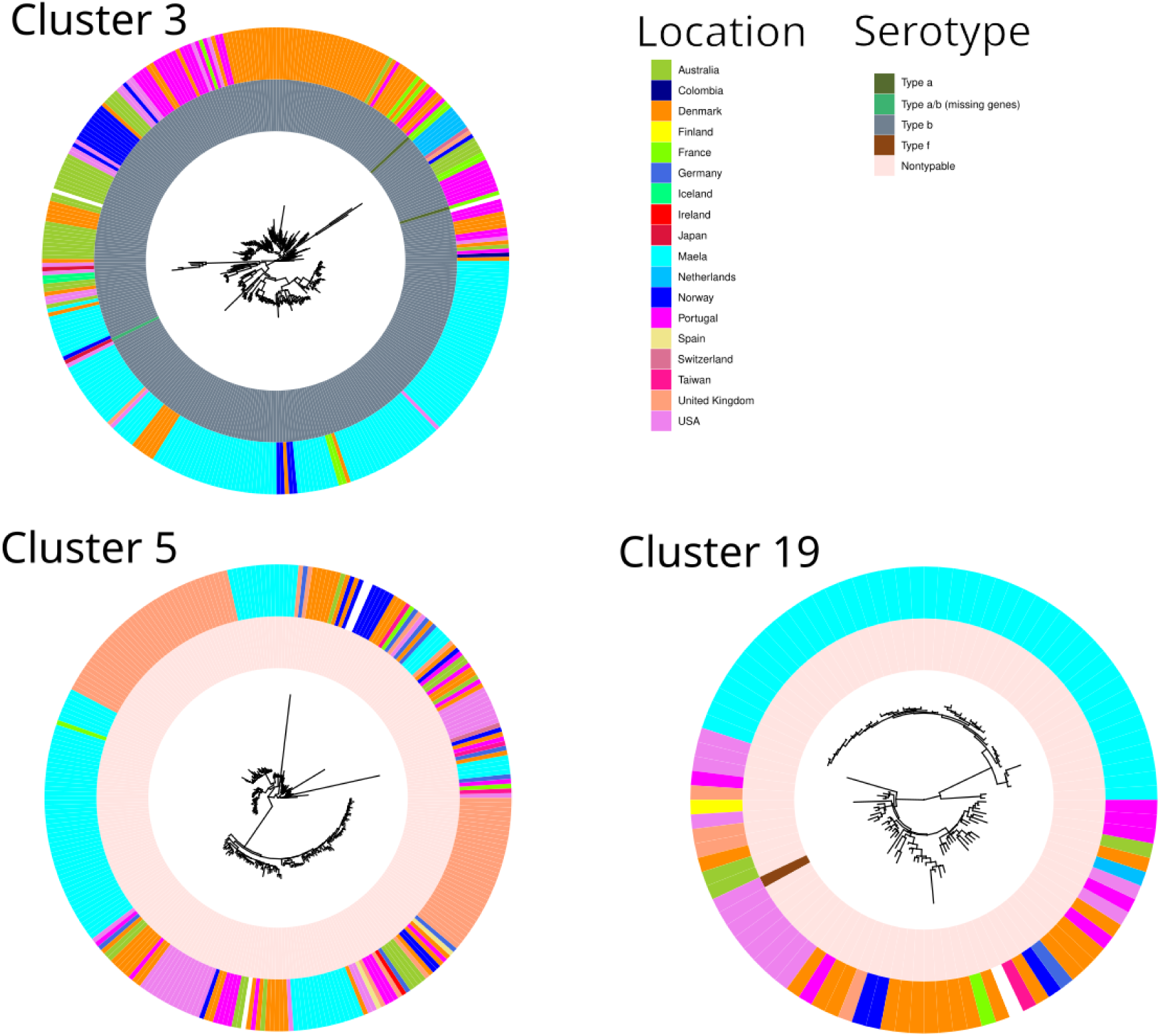
Recombination-free maximum-likelihood phylogenies for each PopPUNK MDR cluster (Clusters 3, 5 and 19), with more than 30% resistance prevalence for a minimum of 4/9 antibiotic classes (see Figure 2), comprising at least 50 genomes. The phylogenies were inferred by using Gubbins on whole-genome pseudoalignments of each cluster, separately mapped against the *H. influenzae* reference 86-028NP (NC_007146.2). *In silico* serotypes (inner ring) and isolation location (second ring) are shown by colour as indicated in the legend. MDR = multidrug resistance. Isolates collected in Maela represent the single-largest origin in all three clusters, at 40% in Cluster 3, 30% in Cluster 5, and 45% in Cluster 19.

## Discussion

The genomic epidemiology of non-b *H. influenzae* has remained largely elusive to date, due to the lack of carriage studies in high-burden settings, particularly in populations from before the rollout of the Hib vaccine. Our study provides the first comprehensive evidence that NT *H. influenzae* are equally capable of causing invasive disease irrespective of their genetic background, even in a pre-Hib vaccine host population. While only colonising isolates were available from the Maela cohort, sampling during episodes of clinical pneumonia has provided insights into disease-associated strains. Results from the multi-country PERCH pneumonia aetiology study ^8^ confirmed a positive association between non-b *H. influenzae* upper respiratory colonisation and chest x-ray confirmed pneumonia. The same study demonstrated an etiologic fraction of 4.5% for non-b *H. influenzae* amongst HIV negative, chest x-ray confirmed cases, which is comparable to the 6.7% fraction estimated for *S. pneumoniae*. There is evidence from several countries of an increasing burden of invasive non-b *H. influenzae* disease, notably in neonates and older adults, with the vast majority being NTHi infections ^1,16^. The high burden of pneumonia attributable to NT *H. influenzae*, and the notable childhood mortality associated with it (3rd most common bacterial pathogen) in the low-resource settings in both Africa and Asia ^16^, combined with the frequent emergence of MDR lineages as identified in the current study, serve as a reminder of the significant health benefits of developing an immunisation program targeting eradication of these pathogens. This would contribute not only towards removing the public health burden of non-B *H. influenzae* invasive disease, but also to significantly reduce both the incidence of AOM and the need to prescribe antibiotics to children.

A pre-vaccine carriage study conducted in The Gambia during the 1980s identified a highly variable carriage rate of serotype b, ranging between 0 and 33% across rural and urban areas, while the species-wide carriage rate was found to be 90% among children under five years of age ^17^. To our knowledge there are no other comparable pre-vaccine studies with serotyping data, but the study in The Gambia suggests that the Maela *H. influenzae* are not atypical in terms of serotype and NT distribution in an unvaccinated host population in a high-burden setting. Post-vaccine carriage studies across Europe and China consistently show a decline of serotype b as expected, but also that NT *H. influenzae* are most commonly colonising young children and that all non-b serotypes are rare ^18–21^. A Belgian study compared carriage rates among children attending day care and those diagnosed with either AOM or invasive disease during 2016-2018. Notably, NT *H. influenzae* were dominating in each category (colonising, AOM, invasive), with the percentages 95.2%, 98.2% and 68.1%, respectively ^19^. Similarly, in Norwegian (2017-2021) and Portuguese (2011-2018) national surveillance of *H. influenzae* invasive disease, NT *H. influenzae* accounted for 71.8% and 79.2% of the cases, respectively. These findings are well aligned with our data from Maela and further with a recent study of community acquired pneumonia in children under five vaccinated against Hib in Vietnam, where a high fraction of NT *H. influenzae* was also detected using real-time PCR ^22^. Interestingly, while serotype b has been found either completely absent ^18,19^ or very rare ^20^ in European carriage studies, it is still found in invasive disease across the continent ^19,23,24^, suggesting ongoing transmission from unvaccinated regions of the world.

Apart from the genomic epidemiology of *H. influenzae*, the overall understanding of the species’ population structure, both encapsulated and NT, has also remained largely elusive, particularly at a global scale, despite various efforts to elucidate it over the past decade. This study, through analysing a large cohort of isolates from an understudied region combined with a systematic collection of publicly available data, suggests that the global *H. influenzae* population is not structured into independently evolving lineages which are predominant in certain regions but rare in others. This is unlike other well-studied bacterial species that colonise the same niche, such as *Streptococcus pneumoniae* or *Neisseria meningitidis,* where distinct, independently evolving lineages have been readily identified for decades ^25–27^. Based on our analyses, *H. influenzae*, in particular NTHi, instead appears to have a population structure reminiscent of panmixia, where routine gene flow between members of the species prevents the formation of stable lineages. This type of population structure would account for the limited success of various methods used to cluster the population in this study, and the difficulties previous efforts have encountered when using smaller datasets ^11^. Technically, clustering methods have likely been limited by the low levels of nucleotide diversity – as low as zero SNPs in over half the core genome of the Maela data – observed within the *H. influenzae* genome, even at a global scale. This, however, is not readily apparent in the output of clustering methods, and only becomes evident when population genomics analyses are conducted at scale.

Despite the low levels of nucleotide diversity, phylogenetic analysis of the combined Maela and globally sequenced isolates remains possible and clearly demonstrates in *H. influenzae* a persistent lack of phylogeographical signal (closely related isolates are highly co-localised), even with this collection of isolates spanning over 50 years. This strongly suggests that inter-regional and intercontinental transmission of these bacteria happens frequently. This is consistent with the high levels of recombination observed in the Maela dataset, as that would facilitate the efficient admixture of migrating isolates with the destination population. Furthermore, the frequent migration and recombination, when combined with widespread evidence of negative selection across coding regions of the genome implied by the low *dN/dS* values, corresponds to a pool of biological forces that likely explains the low nucleotide diversity of the *H. influenzae* genome, particularly the core. It is difficult to disentangle the individual contributions of migration, recombination and negative selection in producing low levels of diversity, and indeed they are likely acting in concert, as has been demonstrated previously in other ecological settings ^28,29^.

These results underscore the importance of a global perspective on disease surveillance when developing public health strategies for managing invasive *H. influenzae* disease, as it is clear that pathogenic adaptations which arise in one part of the world have ample opportunity for global spread. Although the near pan-resistant (six antibiotic classes) MDR lineages we have identified in this study are found more frequently in Maela that any other sampling country, this remains of particular concern as these lineages are all mostly composed of isolates from around the world (55-65%, Fig. 6) and due to the bias of or systematic collection towards high-income settings, we cannot exclude the possibility that these lineages may be further transmitted and established in unsampled LMIC populations with high antibiotic use. Intensified efforts should be made to include *H. influenzae* into AMR surveillance programs as widely as possible. Such surveillance should preferably not be limited to including only bloodstream isolates, because it will otherwise underestimate the prevalence of circulating AMR determinants among pneumonia and AOM clinical cases. Similar to *S. pneumoniae*, carefully conducted studies of *H. influenzae* colonisation in pneumonia cases and controls may provide data on relative invasiveness of capsulated and unencapsulated strains ^30^. Given the significant evidence of adaptation in accessory genes in the Maela population, and that all but one of the pan-resistant MDR lineages was predominantly identified among Maela isolates, it is possible that the camp host population may be exceptionally well-suited to evolutionary adaptation of these bacteria. This could be due to either the host population density resulting in high colonisation and transmission success, or the level of antibiotic use in the camp, and it is further feasible that the fitness cost of maintaining such high levels of resistance beyond these settings is prohibitive. An alternative, and perhaps more likely explanation is that the higher sampling density in the Maela cohort has led to higher statistical power to identify adaptation using methods based on aligned gene sequences, suggesting that similar adaptation could likely have taken place also elsewhere. Widespread genomic surveillance in comparable settings is most likely to lead to the early detection of the spread of extensive levels of AMR, and allow for targeted intervention. Importantly, this type of surveillance data would also be crucial in developing an understanding of the parameters of how selection drives the evolution and maintenance of AMR in *H. influenzae*.

Apart from the concerning implication regarding the possibility of the global spread of AMR in *H. influenzae*, the results of this study also suggest that vaccination may be a particularly effective strategy to control invasive *H. influenzae* disease irrespectively of the serotype, due to the lower level of diversity present within its core genome relative to the accessory, and its highly admixed population structure. Given the low level of observed allelic diversity, the pervasive negative selection we detected throughout the *H. influenzae* genome at a global scale may be strong enough to overcome selection driving compensatory adaptations which would generally reduce vaccine efficacy in response to rollout. Although this is a cause for optimism, it must be tempered by the fact that the high levels of recombination observed in *H. influenzae* may also increase the efficacy of positive selection on any mutations which do arise, as has been observed in other species ^31^. In any case, the stark contrast between the *H. influenzae* population structure identified in this work and the highly stratified population structure of *S. pneumoniae,* both globally ^27^ and in the Maela host population ^32^, strongly suggests that vaccine evasion through inter-lineage competition and replacement, as has repeatedly been observed in *S. pneumoniae* ^33^, would be much less likely to happen in *H. influenzae*, due to the absence of a deeply structured population and local variants. Although there are many complications involved in the design of protein based bacterial vaccines which would need to be overcome^34^, our work supports the conjecture that a single universal vaccine could possibly be developed to combat invasive *H. influenzae* disease, and suggests that the eradication of invasive disease caused by *H. influenzae* may be a feasible end goal of widespread vaccination campaigns. A number of conserved surface antigens have been under investigation as potential candidates for protein subunit vaccines^35^. Recently, antigenic responses to some of the promising candidates have been measured for otitis media -prone children and their controls, these include the recombinant soluble PilA (rsPilA) fused with protein E, protein D and the ubiquitous surface protein A2 (UspA2) from *Moraxella catarrhalis*, as well as ChimV4 (a chimera of protective epitopes from rsPilA), and the outer membrane protein P5 (OMP P5)^36^.

Like other bacteria colonising the upper respiratory tract bacteria and occasionally causing invasive disease, *H. influenzae* has long been known to be naturally transformable and frequently engaging in intraspecific genomic recombination. In this study, we have used two complementary sampling techniques, an in-depth longitudinal sampling of a highly localised population, and a global survey of publicly available data. Population genomic analyses of these data have demonstrated that a high level of recombination, likely acting in concert with negative selection, is important in the evolution of *H. influenzae*. This is primarily through the profound impact on the species’ global population structure, by preventing the formation of stable and independently evolving lineages. This has important implications for how invasive disease caused by *H. influenzae* ought to be controlled, and also raises interesting longer-term evolutionary questions about the underlying genetic drivers of differences between opportunistic pathogenic bacteria colonising the human upper respiratory tract.

## Methods

### Study design and collections

A total of 4,474 *H. influenzae* isolates were retrieved from a mother-infant cohort of 999 pregnant women from the Maela camp for displaced persons, Thailand, from October 2007 to November 2008.^37,38^ Within 24 months’ postpartum period, infants were sampled by nasopharyngeal swabs monthly and when the infant presented symptoms of pneumonia. Of the whole-genome sequences, 3,970 passed the quality control and were included in the genomic analyses (appendix 1 p 1). For comparative analyses, a systematic search was conducted for publicly available short-read genome sequences for which country and year of isolation metadata was available. 6129 isolate data were retrieved from the ENA ^9,10,13,39–66^, of which 5,879 passed quality control, resulting in a final dataset size of 9,849 isolates.

### Sampling and Sequencing Procedures

Between October 2007 and November 2008, 999 pregnant women from the Maela camp for displaced persons (located on the Thailand-Myanmar border in Tak province, NW Thailand, Fig. 6a) were recruited into a mother-infant colonisation study. Infants were followed from birth for 24 months and a nasopharyngeal swab specimen collected (dacron tipped swabs; Medical Wire & Equipment, Corsham, UK) at monthly intervals and if the infant presented to the Shoklo Malaria Research Unit clinic with symptoms and signs compatible with WHO clinical pneumonia (Fig 6b).

Following sampling, the nasopharyngeal swab (NPS) tip was excised immediately into a sterile cryovial containing 1mL STGG (skim milk, tryptone, glucose, glycerol medium; prepared in-house) using 70% ethanol-cleaned scissors. NPS-STGG specimens were transferred to the SMRU microbiology laboratory in a cool box, within eight hours of collection, and were frozen at −80°C until culture.

Ten microlitres of thawed NPS-STGG specimen was cultured onto plain chocolate agar (Clinical Diagnostics, Bangkok, Thailand), a 10 unit bacitracin disc (Oxoid, Basingstoke, UK) applied to the first streak, and the plate incubated overnight at 36°C in 5% CO_2_. Bacitracin resistant colonies were confirmed as *H. influenzae* by Gram stain and X+V factor dependent growth. Serotype was determined by slide agglutination (Becton Dickinson, Franklin Lakes NJ, USA). Pure isolates of *H. influenzae* were harvested from an overnight culture plate into 1mL STGG and stored at −80°C prior to DNA extraction and sequencing.

Short read whole-genome sequencing of the 4,474 *H. influenzae* isolates was performed at the Sanger Wellcome Institute on Illumina-HTP NovaSeq 6000 platform with 150bp paired-end sequencing (appendix 1 p 1).

For long-read sequencing, one reference isolate was selected per each of the 48 largest PopPUNK clusters, covering 3,558 (89·6%) of 3,970 isolates of the study cohort. Reference isolates were selected based on the gene presence absence matrix from the estimated pangenome, using a published selection pipeline (https://gitlab.com/sirarredondo/long_read_selection).^67^.

The selected *H. influenzae* strains were subcultured on chocolate agar and incubated overnight at 35-37°C in 5% CO2. Genomic DNA was extracted using the Qiagen MagAttract HMW DNA Kit. Whole-genome sequencing libraries were constructed using the Oxford Nanopore Technologies SQK-NBD112.96 Native Barcoding Kit and all 48 strains were pooled together and sequenced on one ONT R9.4.1 flowcell using a MinION Mk1c. Hybrid assembly of the reference isolates was performed using a publicly available pipeline (https://github.com/arredondo23/hybrid_assembly_slurm) with minimum ONT coverage of 40x and a phred score of 20 to trim the Illumina reads, resulting in 40 (83·3%) complete hybrid assemblies of the 48 reference isolates.

### Genomic Analysis

A total of 4,474 Haemophilus influenzae isolates were sequenced at the Sanger Wellcome Institute (Hinxton, UK) on the Illumina-HTP NovaSeq 6000 150 basepair paired-end platform. Species contamination was identified by using Kraken v.0.10.6.^68^, and the sequence data failed quality control if the depth of coverage was <20x, or if there was evidence of contamination or mixed strains, poor assembly, or extreme violation of any of the quality control parameters. Short-read genome sequences, both newly sequenced and publicly available, were assembled and annotated using a published pipeline with default parameters,^69^ and a round of QC on all isolates was performed based on the number of contigs, genes, and distance from the origin in an MDS projection of all pairwise distances. Isolates were clustered using PopPUNK v.2.4.0 ^70^ separately on the Maela data, with core threshold of 0.11, and on the combined global data with default thresholds. Antimicrobial resistance genes and point mutations were screened from assemblies using AMRFinderPlus v.4.0.3 and *H. influenzae*-curated database v.2024-12-18.1 (--organism Haemophilus_influenzae) with minimum identity of 75% and minimum coverage of 80%. Hicap v.1.0.3 ^12^ was used to infer capsule type from assemblies. Core Genome Multi-locus Sequence type (cgMLST) was identified from each isolate’s pan-genome genes using chewBBACA ^71^ and the *H. influenzae* cgMLST database^11^, and a simple network clustering method was used to group isolates into complexes based on the number of mismatches in their allele profiles, of either 100 or 250 allelic mismatches. For phylogenetic analyses on the Maela data, the sequence reads of the 3,970 quality control-passed genomes were mapped to the complete genome of *H. influenzae* 86-028NP (NC_007146.2)^72^ using Snippy v.4.6.0 (https://github.com/tseemann/snippy) and a SNP-only sequence alignment was created using snp-sites v.2.5.1.^73^ A phylogeny for Maela collection was inferred using FastTree v.2.1.10 with a generalised time-reversible model ^74^, and a maximum likelihood tree was inferred using IQ-TREE v.2.4.0^75,76^ on Panaroo core-genome alignment, with uninformative regions masked using information entropy scores, to analyse the combined global collection.

The pangenome was inferred for the Maela genome collection using Panaroo v.1.2.9 ^77^ using sensitive mode and merging paralogs. The pangenome was further inferred for the entire combined global dataset running Panaroo in strict mode. FastGEAR v.2016-12-16 was used to infer recombinations, and pixy v.1.2.7.beta1 was used to infer per-gene nucleotide diversity, π, both from aligned pan-genome gene sequences. Genomegamap v.1.0.1 ^78^ was used to infer maximum-likelihood estimates of each aligned gene average *d*N/*d*S for each gene in the pangenome of the entire collection. In keeping with the method as described, only genes with an estimated nucleotide diversity (theta) value greater than 0.005 were considered robust enough estimates for further interpretation, leading to successful estimates for only 6,853 genes. Genes with robust estimates of *dN/dS* greater than 2 were further analysed using the HYPHY package v.2.5.60 ^79^, specifically the FUBAR, FEL, and ABSREL statistical tests for pervasive gene-wide directional selection, specific sites subject to directional selection, and subsets of branches subject to directional selection respectively. Genes which had significant results to at least one of these HYPHY tests were manually investigated by searching the consensus and variant nucleotide sequences against the non-redundant protein database with tblastx, searching ESMFold-predicted protein structures against the AlphaFold, Uniprot, and Swiss-prot database using Foldseek. 3D structural alignments of consensus, variant, and reference protein structures were created for specific genes using TM-align.

Multidrug resistant (MDR) clusters in the combined collection were defined as PopPUNK clusters with a minimum of 50 isolates per cluster, of which at least 30% harboured resistance determinants to at least 4 of 9 antibiotic classes (aminoglycoside, β-lactam, phenicol, sulphonamide, tetracycline, trimethoprim, macrolide, quinolone, rifamycin). For phylogenetic reconstruction of the global MDR clusters, assemblies of each cluster were mapped to the reference *H. influenzae* 86-028NP ^72^ using Snippy and phylogenies inferred using Gubbins ^80^.

In order to test for an association between lineages of NT isolates and disease, we first de-duplicated isolates from the same host and likely to be clonally related, to remove the bias associated with longitudinal sampling. To do this, we grouped all isolates collected from the same hosts within 60 days belonging to the same PopPUNK cluster, and randomly selected a single isolate to keep in the analysis, while the rest were excluded. Then, for all remaining NT isolates spanning 120 PopPUNK clusters in the Maela cohort, we performed a permutation test of association between genetic lineages and pneumonia, by randomly shuffling “Status” labels (carriage and pneumonia). A two-sided Fisher’s exact test with 10,000 Monte Carlo replications was used.

### Ethical approval

Written informed consent was obtained from the participating infants’ mothers prior to enrolment into the cohort study. Ethical approval was granted by the ethics committees of the Faculty of Tropical Medicine, Mahidol University, Thailand (MUTM-2009-306) and Oxford University, UK (OXTREC-031-06). The sequencing work on stored isolates described here was approved by the same committees (TMEC-19-043; OxTREC-551-19).

### Reporting summary

Further information on research design is available in the Nature Portfolio Reporting Summary linked to this article.

## Data availability

All sequence data generated as part of this work is available on the ENA/SRA/DDBJ under study accession PRJEB41043. Metadata for both newly sequenced data and the systematic global collection is available on microreact at the links as indicated in figure captions. (https://microreact.org/project/oMm8PFCoG2429JwiDBpdru-maela-h-influenzae and https://microreact.org/project/ioyt4oJRSJgeFGK9KmFyVk-global-h-influenzae-core-tree) Sequence data was produced according to Illumina protocols, and publicly available data was downloaded from the ENA using enadownloader. No novel algorithms or computational methods were developed during the course of this work.

## Acknowledgements

This work was supported by European Research Council (grant 742157; to JC), Wellcome Trust Grants (206194, 083735; to PT, MORU core award 220211; to PT; 206194, 220540/Z/20/A to Wellcome Sanger Institute), Trond Mohn Foundation (BATTALION grant; to JC, AKP, SM, and NM), Marie Skłodowska-Curie Actions (Grant 801133; to AKP, SM, and NM).

## Author information

These authors contributed equally: Neil MacAlasdair, Anna K. Pöntinen, Clare Ling. These authors jointly supervised the study: Paul Turner, Jukka Corander

### Authors and Affiliations

Department of Biostatistics, University of Oslo, Oslo, Norway

Neil MacAlasdair, Anna K. Pöntinen, Sudaraka Mallawaarachchi, Jukka Corander

Parasites and Microbes, Wellcome Sanger Institute, Hinxton, UK

Neil MacAlasdair, Stephen D. Bentley, Jukka Corander

Cambodia Oxford Medical Research Unit, Angkor Hospital for Children, Siem Reap, Cambodia

Clare Ling, Claudia Turner, Paul Turner

4 Centre for Tropical Medicine and Global Health, Nuffield Department of Medicine, University of Oxford, Oxford, UK

Clare Ling, Francois H. Nosten, Claudia Turner, Paul Turner

Peter MacCallum Cancer Centre, Melbourne, Victoria 3052, Australia

Sudaraka Mallawaarachchi

Sir Peter MacCallum Department of Oncology, University of Melbourne, Melbourne, Victoria 3052, Australia

Sudaraka Mallawaarachchi

Mahidol-Oxford Tropical Medicine Research Unit, Faculty of Tropical Medicine, Mahidol University, Bangkok, Thailand

Janjira Thaipadungpanit

Shoklo Malaria Research Unit, Mahidol-Oxford Tropical Medicine Research Unit, Faculty of Tropical Medicine, Mahidol University, Mae Sot, Thailand

Francois H. Nosten

MRC Centre for Global Infectious Disease Analysis, Department of Infectious Disease Epidemiology, Imperial College London, London, UK

Nicholas J. Croucher

Department of Mathematics and Statistics, University of Helsinki, Helsinki, Finland Jukka Corander

## Contributions

N.M. and A.K.P. had the major responsibility in bioinformatics, population genomics and statistical analyses. C.L. Management and provision of swab specimens and isolates, curation of isolate data. S.M. conducted additional statistical analyses. J.T. Investigation of isolates and MinION sequencing. F.H.N. Administered the cohort study. C.T. Investigated and curated cohort clinical data. S.D.B. advised on study design and interpretation of results. N.J.C. advised on population genomics and interpretation of results. P.T. and J.C. acquired funding and jointly designed and supervised the study. N.M., A.K.P., P.T. and J.C. jointly wrote the initial draft and all other authors improved the paper.

## Additional information

Extended data

Extended Data Fig. 1.

Unnamed Gluconate Transporter (pangenome ID: group_3361) Identified as being under selection in some Maela isolates

A: Structural alignment of variant under selection in Maela data (Blue) and wild type variant (Gold), with the large domain composed of alpha helices at the C-terminus deleted in the Maela variant. B: Interpro scan domain annotation of the wild-type protein indicating that the deleted C-terminus domain in the Maela data is transmembrane

Extended Data Fig 2.

BrnT toxin (pangenome ID: group_4059) variant includes an altered alpha helix domain A: Structural alignment of the structural variant found in Maela isolates (blue) compared with the wild type *H. influenzae* structure (gold). The variant under selection contains some amino acid changes, as well as a small deletion, leading to the absence of an alpha helix when compared to the wild type *H. influenzae* structure. B: Structural alignment of both *H. influenzae* BrnT genes compared to the reference structure from *Brucella abortus*.

## Supplementary information

Supplementary Table 1. Table of the main PopPUNK clusters, the number of isolates, predominant serotype, cgMLST, and Mismatch 100 cgMLST complex in each cluster. Percentages of the predominant serotype, cgMLST, and 100 mismatch cgMLST complex are provided in parentheses.

Supplementary Table 2. Sequencing QC information for all newly sequenced isolates in this study, including those which failed QC

Supplementary Table 3. Contamination QC information for all the combined collection of the systematic global collection of all publicly available isolates and the newly sequenced Maela collection. QC metrics include the number of genes annotated by Prokka, the number of contigs in the assembled genome, and the distance from the origin of the MDS projection of all pairwise distances as calculated by mash.

